# Categorical rhythmic priors in macaques

**DOI:** 10.64898/2025.12.29.696861

**Authors:** Ameyaltzin Castillo-Almazán, Luis Prado, Nori Jacoby, Hugo Merchant

## Abstract

Rhythmic ability is a universal aspect of human cultures and sets the basis for musical rhythm perception and synchronization. While humans can synchronize movements to complex rhythms, it is unclear whether this capability extends to our primate ancestors. In this study, we explore whether primates can synchronize to complex rhythms and acquire rhythmic representations with generalizability akin to humans. Using controlled behavioral experiments, we provide evidence that monkeys not only can synchronize to short-long or long-short rhythms but also learn representations that generalize across a wide range of rhythm ratios and total durations. These results indicate ability to flexibly represent ratios within a relative timing framework is not exclusive to humans, but it is also present in monkeys. In addition, the produced intervals show a bias towards rhythmic categories. Notably, in an iterative tapping task, macaques and humans showed large priors for isochrony and integer ratios (2:1, 3:1). These results demonstrate a common biological foundation for rhythm synchronization in primates, extending our understanding of the shared cognitive mechanisms between primates and humans, and highlight the enormous potential of using monkeys to study the neurophysiological basis of complex rhythm perception.

## Introduction

Music cognition is one of the traits that make us human(Honing et al., 2012; Merchant, Grahn, et al., 2015). The rhythmic structure of music follows at least three universals across all human cultures, namely, the perception of an isochronous (regularly periodic) beat, the hierarchical organization of rhythms of unequal strength (defining strong and weak beats), and the bias for grouping rhythms in two (e.i. a march) or in three (e.i. a waltz)(Ravignani & Madison, 2017; Savage et al., 2015). Furthermore, the preference for specific ratios like 1:2 and 1:3 is maintained for a wide range of tempi, supporting the notion of relative rhythmic timing, with these ratios as prototypes or categories. Strong evidence for this ratio preference in humans comes from a recent method that uses the iterated reproduction of random sequences to estimate the categorical bias or internal priors of rhythms(Jacoby & McDermott, 2017). In this paradigm, listeners are first exposed to a sequence of sounds separated by random seed intervals and are instructed to reproduce the sequence by tapping in synchrony with the rhythmic stimuli. Then, the original sequence of stimuli is replaced by the average intervals reproduced by the listener. It has been shown that the listeners reproduction depends on a Bayesian process where performance depends on the integration of the currently heard rhythm with a prior. Hence, across iterations the production biases emerge and converge towards the listeneŕs priors. Two– or three-interval rhythms have been used in these experiments, with a uniform seed distribution, covering a wide range of ratios and total durations. Notably, humans show mental rhythmic categories at the isochrony (1:1, 1:1:1) and small integer ratios, such as 1:2 or 1:1:2. These rhythmic categories are present for different total durations, confirming a relative representation of rhythm. In addition, a recent large scale cross-cultural study using the same iterative tapping task, demonstrated that although the categorical small integer ratios are present across cultures there is also a substantial influence of the local musical environment(Jacoby et al., 2024)

In this context, a critical question is what are the evolutionary origins of rhythm perception and synchronization? An influential hypothesis in the field is the vocal learning and rhythmic synchronization hypothesis, which states that high vocal learning is a preadaptation for the spontaneous perception and synchronization to musical beat(Patel, 2010, 2021). This idea is supported by the observation that most musical species do tend to also have complex vocal communication, such as humans and songbirds(Patel et al., 2009). Nevertheless, this hypothesis also predicts that apes and monkeys, which are modest vocal learners, cannot properly perceive and entrain to rhythms. This notion has been strongly challenged recently. On the one hand, EEG studies in the Rhesus monkey have shown that macaques produce evoked potentials linked to the detection of isochronous auditory patterns(Honing et al., 2018), as well as to subjectively accented 1:2 and 1:3 rhythms from auditory metronomes(Ayala et al., 2017; Criscuolo et al., 2023). On the other hand, biomusicologists have shown that great apes show rhythmic drumming behaviors(Ravignani et al., 2013) and can learn to execute human-like sequences(Dufour et al., 2015; Fuhrmann et al., 2014), while lemurs produce in the wild coordinated songs with isochronous and 1:2 rhythmic categories(De Gregorio et al., 2021). Finally, monkeys trained on tapping tasks can flexibly and predictively produce periodic intervals in synchrony with auditory and visual metronomes(Betancourt et al., 2023; Gámez et al., 2018), can continue tapping without sensory cues(Zarco et al., 2009), and can even consistently tap to the subjective beat of different music excerpts(Rajendran et al., 2025). Hence, these studies suggest that non-human primates can perceive and synchronize to simple rhythms such as isochrony. This implies that they have the brain sensorimotor machinery to: (1) extract a rhythm from a continuous stream of sensory events, (2) generate an internal rhythmic signal that predicts the future beat events, and (3) produce anticipatory motor commands such that movements coincide or slightly anticipate the next rhythm(Lenc et al., 2020; Merchant & Honing, 2014).

A large set of functional imaging studies have shown that the neural substrate of the human internal rhythmic clock lies in the voluntary skeletomotor system, including the medial premotor areas (MPC: SMA and pre-SMA), the basal ganglia (most often the putamen, but also caudate nucleus and globus pallidus) and the motor thalamus, which form an important cortico-basal ganglia loop(Grahn & Rowe, 2009; Kung et al., 2013; Sánchez-Moncada et al., 2024). However, studies of rhythmic timing using non-invasive techniques lack the spatial and temporal resolution necessary to determine the neural dynamics and interareal interactions underlying perception and synchronization to rhythms. Due to their rhythmic abilities and evolutionary close audiomotor system, Rhesus monkeys are key animal models for understanding the neural mechanisms behind rhythm perception and synchronization. In fact, recent neurophysiological studies in monkeys indicate that the internal pulse representation during rhythmic tapping to visual and auditory metronomes depends on the neural population dynamics in MPC. A key property of MPC neurons is the relative representation of beat timing. Cells that encode elapsed or remaining time for a tap show up-down ramping profiles that span the produced interval, scaling in speed as a function of beat(Merchant, Zarco, et al., 2011; Merchant & Averbeck, 2017). In addition, these cells are recruited in rapid succession producing a progressive neural pattern of activation that flexibly fills the beat duration depending on the tapping tempo, providing a relative representation of how far an interval has evolved(Crowe et al., 2014; Mello et al., 2015; Merchant, Pérez, et al., 2015; Zhou et al., 2020). The neural cyclic evolution is more evident when the time-varying activity of MPC neurons is projected into a low-dimensional state space(Gámez et al., 2019). The population neural trajectories show the following properties. First, they have circular dynamics that form a regenerating loop for every produced interval. Second, they converge in similar state space at tapping times, resetting the beat-based clock at this point. This internal representation of rhythm could be transmitted as a phasic top-down predictive signal to sensory areas(Lenc et al., 2020). Third, the periodic trajectories increase in amplitude and decrease in speed as a function of the length of the isochronous beat encoding the tempo of the metronome(Betancourt et al., 2023; Gámez et al., 2019). Finally, using auditory and visual metronomes it is evident that there is an amodal internal representation of tempo in the amplitude and speed of the MPC neural trajectories that interacts with a modality specific constant external input^18^. These functional properties indicate an underlying prerequisite for invariant representation of rhythmic ratios, as MPC neurons can encode duration relative to a reference. Could these features manifest in monkey behavior? Specifically, could a monkey synchronize with a complex rhythm across a broad range of tempi?

Here we trained macaques in a two-interval rhythm tapping task to test whether monkeys can produce short-long and long-short rhythms and then determine the possible rhythmic priors in non-human primates thanks to the iterated task. We found that monkeys can reproduce a sequence of multiple two-interval rhythms, and they can flexibly change the produced intervals depending on cued rhythms with different ratios and total durations. In addition, monkeys present a large prior for isochrony, as well as for some integer (2:1, 3:1) and non-integer ratios (0.8:2) in the iterated task. These findings support the notion that monkeys share with the human two key abilities for music cognition: complex rhythm perception and execution, and the inherent bias towards rhythmic categories. This also suggests that, like humans, monkeys may possess a categorical representation that remains invariant across different tempi.

## Results

### Predictive and malleable tap isochrony

We trained two monkeys in a visual Isochronous Synchronization Tapping Task (ISTT, see Methods). The animals sat in a primate chair and started a trial by holding a lever during the presentation of a yellow background with the first flashing red-square that acted as a metronome presented on a LED matrix (Fig. 1a). The disappearance of the yellow background was the go-signal to tap a button seven times in synchrony with the red-square metronome, generating seven inter-response intervals (IRI0 to IRI6). The animals received a liquid reward if their accuracy and asynchronies were below an error threshold (see Methods). We used 16 inter-stimulus intervals (ISI) (400 – 1,150 ms), which were presented pseudorandomly in blocks of five training trials, during which the monkey adapted to the new tempo, and 20 testing correct trials.

**Figure 1.**
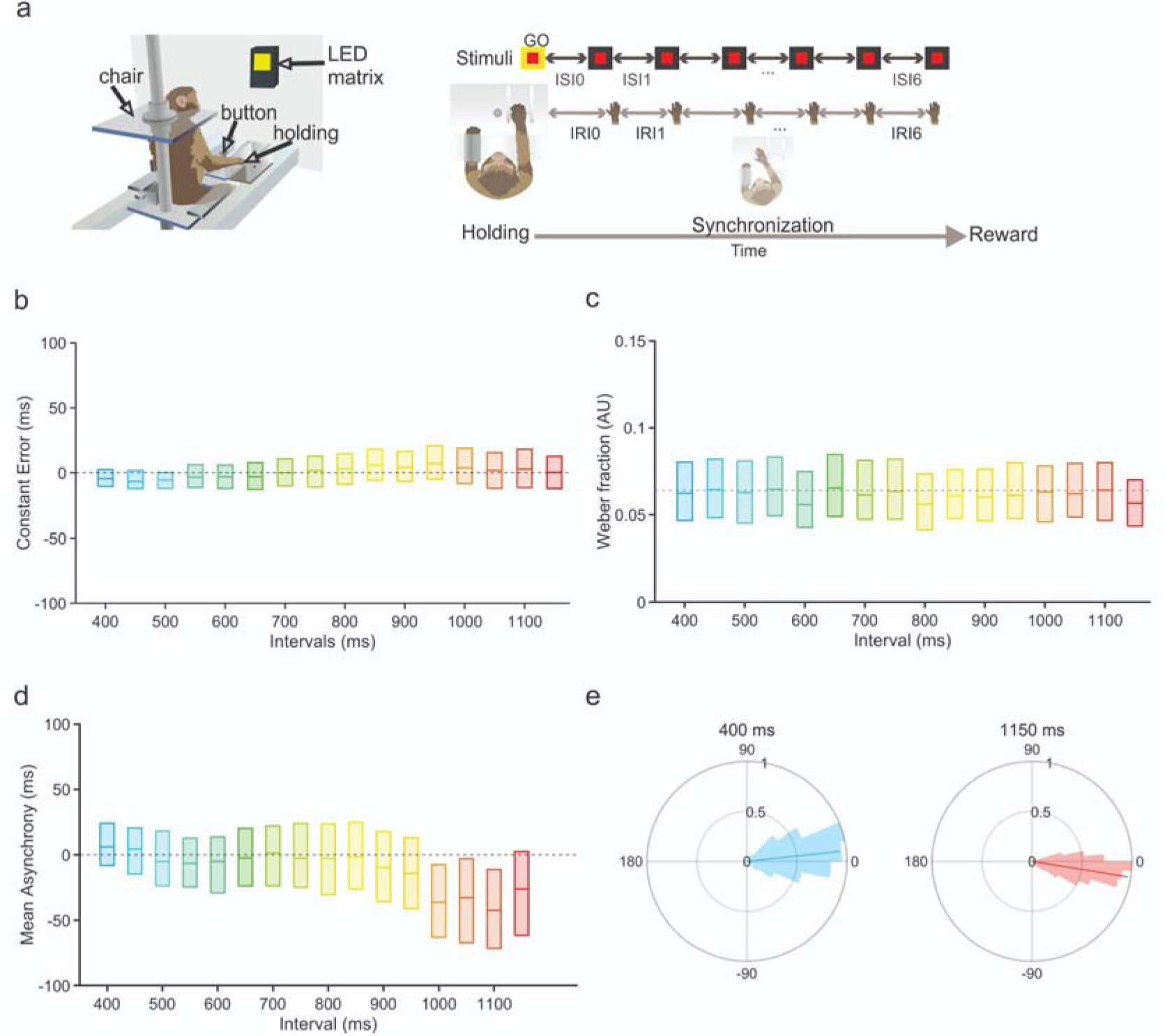
Isochronous tapping synchronization task and behavior for monkey M. **a**, Experimental setup and trial structure. A trial started when the monkey held a lever during the presentation of a yellow background and the first flashing red-square that acted as a metronome. The disappearance of the yellow background was the go-signal to tap a button seven times in synchrony with the red-square metronome, generating seven inter-response intervals (IRI0 to IRI6). The first produced interval (IRI0) was not analyzed since it included the movement from the lever to the button. The inter-stimulus interval (ISI) of the metronome was randomly selected from 16 intervals (400 – 1,150 ms). The monkeys performed five training and 20 testing correct trials in one interval before changing pseudorandomly to a block of trials with another interval. **b**, Boxplots of the Constant Error as a function of interval showing the median (central line), the 25th (bottom edge), and 75th percentiles (top edge). **c,** Boxplots of the Weber fractions across tempos. **d**, Asynchrony Boxplots. The monkey’s taps were predictive, with asynchronies close to zero across the wide range of tested intervals. **e**, Polar plots showing examples of the distributions of the asynchronies for the extreme tested intervals: shorter 400 ms and longer 1150 ms. The mean resultant was close to one (shown as a vector; 0.93 and 0.97, respectively, Rayleigh test p<0.0001) (see Figure S1 for all polar plots for both monkeys).

We used three measures of performance: the constant error is the difference between the produced IRI and target ISI (Figure 1b, Figure S2a), the Weber fraction is the standard deviation of the IRIs divided by the ISI (Figure 1c, Figure S2b), the asynchrony corresponds to the time difference between the stimulus onset and tap onset (Figure 1d, Figure S2c). All three measures showed tempo invariance, with small main effects of tempo in the one-way ANOVA (F(15,6399) = 25.33, p < 0.0001 η^2^ = 0.056; F(15,6399) = 5.5, p< 0.0001 η^2^ = 0.013; F(15,6399) = 77.15, p < 0.0001 η^2^ =0.15, for constant error, Weber fraction, and mean asynchrony, respectively). Despite the wide range of used tempos, the monkeys flexibly and accurately changed their produced durations and showed a similar Weber fraction (Figure 1b,c). Notably, for each tempo, the distribution of asynchronies in polar coordinates showed a consistent phase that was unimodal (significant Rayleigh’s test, p < 0.0001), and where the corresponding mean resultant had an angle close to zero and a magnitude close to unity (Figure 1e, Figure S1). This strong predictive behavior also showed evidence of a preferred tempo around 650-850ms, since the asynchronies were not different from zero in this range (t-test (15,6399) p = 0.14934, 0.91919, 0.1813, 0.024902, 0.88651 with Bonferroni correction for multiple comparisons). Therefore, the tapping synchronization of monkeys to visual metronomes showed three hallmark properties of human rhythmic behavior: the accuracy and Weber fraction of the produced intervals are tempo invariant, the tapping is predictive, and the existence of a preferred tempo(Repp, 2005; Repp & Su, 2013).

### Complex rhythms reproduction

To our knowledge, the ability to reproduce complex rhythms has never been demonstrated in monkeys. We used a visual version of the two-interval rhythm task (2iRT), previously used in humans(Jacoby & McDermott, 2017), to determine whether macaques could reproduce short-long (first interval shorter than second) or long-short rhythms (first interval longer than second; Figure 2a, middle). Rhythms can be described in terms of ratios corresponding to the proportion of the first interval by the total pattern duration (Figure 2a, bottom). For example, musical rhythms across human cultures tend to have a simple integer ratio of 1:2 (or a ratio of 0.33) (Figure 2a, bottom). After the training in the isochronous task the monkeys were tested on the 2iRT, where they produced three repetitions of the two-interval rhythm for a total of six taps. Remarkably, since the first day that the monkeys were confronted with the 2iRT, they were capable to systematically produce intervals of different duration (Weber fraction first day: interval 1 = 0.050 CI=[0.017, 0.093]; interval 2 = 0.049 CI=[0.008, 0.104]), suggesting that macaques have an inherent rhythmic clock that can time complex ratios not only isochrony (Figure S3). However, in these days the monkeys reversed the short-long pattern with a long-short pattern behavior and vice versa, probably because they were reproducing the previously presented interval in the sequence (Figure S3). Hence, to give more information to the animals we used during training small green squares for short intervals and large red squares for long intervals (Figure 2a). In addition, to train the monkeys without biasing their behavior for specific ratios, we tested the macaques in the 2iRT using 53 ratios (from 0.29 to 0.71) and eight total durations (1,000 to 1,700 ms), for a total of 424 rhythm combinations that were presented pseudorandomly during more than two months of training. Notably, monkeys could reproduce properly a wide variety of rhythmic patterns (see Figure 2b, Figure S4a for exemplary sessions). Quantitatively, they displayed a low Weber fraction with a mean = 0.069; with small but significant variation with the total duration of the rhythm (F(7,415), p = 1.19e-29; see Figure 2c). The mean asynchrony was slightly positive for short and slightly negative for long total durations (M=11.53 ms CI=[-38.85 65.66]), with values not different from zero for 1500 and 1600 ms (p> 0.5 via t-test), supporting the notion of a robust predictive behavior across a large span of two-interval rhythm durations. In addition, the produced intervals 1 and 2 of the 2iRT changed systematically as a function of the target ratio and the total duration of the rhythm (Figure 2e, separate ANOVAs for the two intervals using total duration and ratio as factors, all main effects and interactions P < 0.0001), suggesting a highly flexible ability to reproduce two-interval rhythms across a wide range of total durations (see also Figure S4b). As a complement, monkeys show evidence for ratio internalization. The produced ratios showed small modulations by the total duration (F(7, 4991)=11.66 p=8.96e-15 η^2^= 0.0012) and followed a categorical behavior with larger produced ratios for small target ratios and shorter produces ratios for long target rations with a categorical behavior (Figure 2f left; slope of shorter and longer ratios was alpha= 0.32 and alpha= 0.27, respectively, which were significantly different from a slope of 1). This phenomenon is also evident in humans (slope of shorter intervals and longer was alpha= 0.48 and alpha= 0.53; Figure 2f right) and is consistent with human categorical effects extensively studied in previous literature(Desain & Honing, 2003; Repp et al., 2012; Roeske et al., 2020). Therefore, these findings suggest that monkeys learned a tempo invariant representation of ratios, representing timing in a relative rather than in absolute fashion, consistent with the evidence of relative timing in the neural representation of isochronous timing of medial premotor areas in the same species^18^.

**Figure 2.**
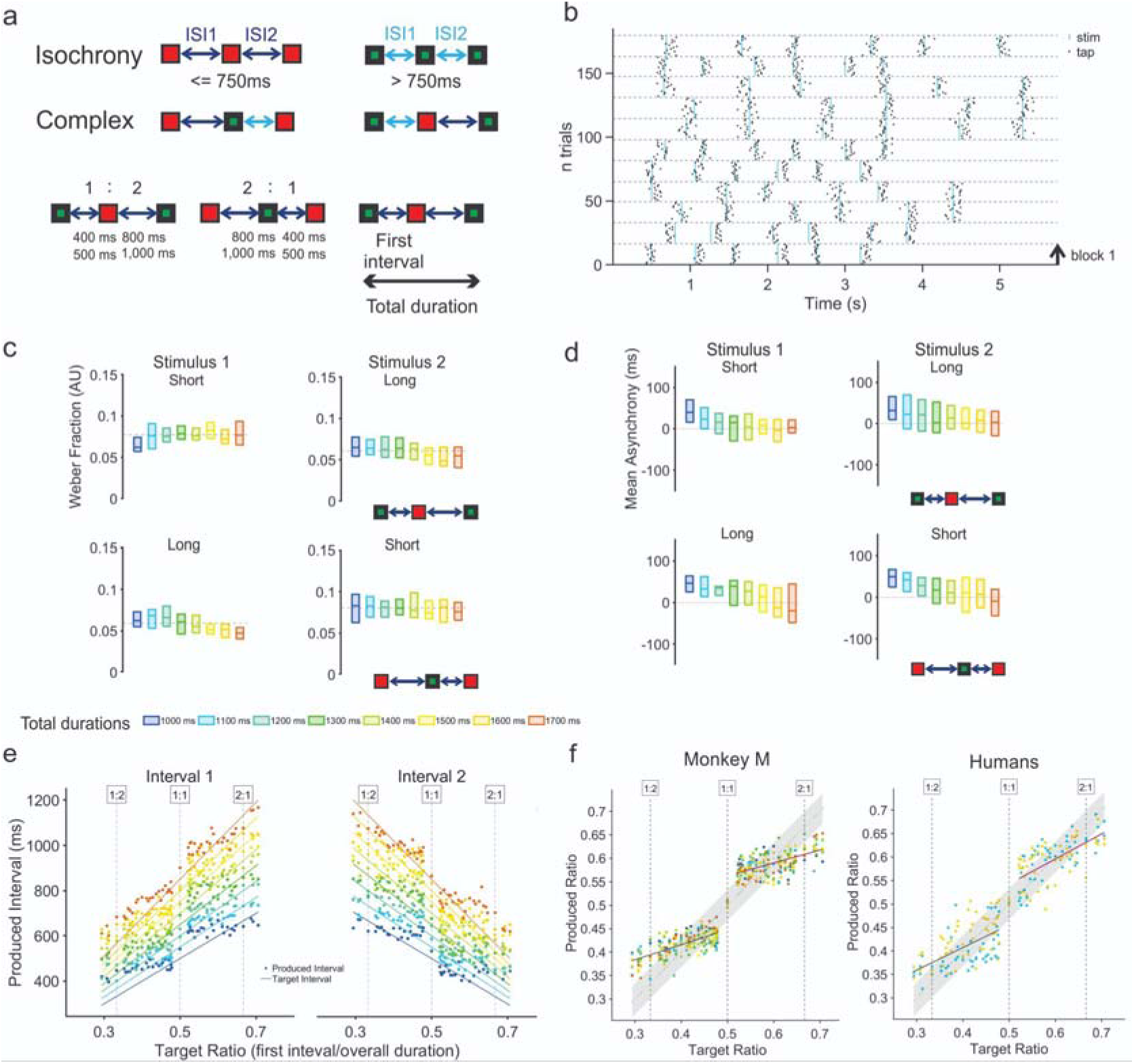
Two-interval rhythm synchronization task and behavior. **a,** Different configurations for the two-interval rhythms, we distinguish the interval duration with a specific color: for intervals bigger or equal to 750 ms, large red squares; for intervals smaller than 750 ms, small green squares. We tested isochronous rhythms, top, and complex rhythms (long-short or short-long), middle. Examples of two different ratios, 1:2 and 2:1, bottom, the first interval of the rhythm is a proportion of the second interval (for 1:2, first interval = 500 ms x 1 then the second interval = 500 ms x 2 = 1000 ms). **b,** Raster plot for a session of the two-interval rhythm task. Each block has 20 trials of the same interval ratio and total duration combination. The horizontal dashed gray line delimits each block. Black dots correspond to taps and blue lines to each presented stimulus. Note the close relation between tapping times to each stimulus of the rhythm. **c,** Weber fraction for each stimulus of the two-interval rhythm for all the tested total durations (the 8 total durations are color-coded below). The mean Weber fraction is marked by a gray dashed line, and each boxplot presents the median (central line), the 25th (bottom edge), and 75th percentiles (top edge). **d,** Mean asynchrony boxplot for each stimulus of the two-interval rhythm for all the tested total durations. **e,** Mean produced intervals, for the first and second interval of the 2iRT, as a function of the 53 target ratios for each of the 8 total durations. The intervals were grouped on short-long (interval 1, between 0.29 and <0.5) or long-short ratios (interval 2, between >0.5 and 0.71). Dashed gray lines indicate integer ratios on the top. **f,** Mean produced ratio as a function of the target ratio for each total duration for monkey M and humans. Each colored point corresponds to the produced ratio. The gray shaded area represents the range of the produced ratio to be considered near the target ratio (see methods). The black and magenta lines correspond to a linear regression for the mean produced ratio for the set of 8 total durations in the configuration shot-long or long-short respectively. Panels d-f have the same color code as c.

### Wider range of ratios

The large flexibility of beat performance inspired our exploration over a wider range of ratios, equivalent to those tested in humans. This task was called the full-range 2iRT since we used 54 ratios from a uniform distribution ranging from 0.18 to 0.82. We chose the total duration of 1600 ms, where monkeys were very proficient. Crucially, in this case, the metronome was a uniform flashing square; hence, no short/long cues were provided to the monkey. As observed from the previous experiment, the Weber fraction was small (X = 0.077 CI=[0.011 0.254], Figure S5a), the mean asynchrony was not significantly different from zero (X = –0.622 CI=[-61.31, 69.42], Figure S5b), and produced intervals had a positive slope (produced interval 1 against target interval: alpha = 0.94 p<0.01, produced interval 2 against target interval: alpha = 0.95, p<0.01). An analysis of two-way ANOVA for each produced interval shows that the interaction between the ratio and the serial order of the interval within the rhythm is statistically significant for the produced interval, meaning that the monkey can differentiate and combine two different interval durations in a pattern (Interval 1: F(26, 1527) = 168.31, p < 0; Interval 2: F(26, 1527) = 149.35, p < 0). Remarkably, the produced ratio was very similar to the target ratio across the wide tested range of ratios (slope 0.92, p<0.00001). The corresponding ANOVA shows a significant difference between the target ratios and the interval configuration, as well as the corresponding interaction, p < 0. These results confirm that the monkey has an internal representation of relative timing for a quite wide range of ratios.

### The rhythm priors revealed by iterated reproduction

As commented above, humans and monkeys present a categorical effect when synchronizing to a two-interval rhythm (Figure 2f). To assess the perceptual prior underlying the categorical effect on the 2iRT in humans and monkeys, we used a previously used iterated task^5^ using visual stimuli for the monkey and visual and auditory stimuli for humans. Critically, it has been shown previously that the execution of this type of task enables the amplification of biases and the identification of preferred categories (Griffiths et al., 2024; Jacoby et al., 2024; Jacoby & McDermott, 2017). In our version of the task, each trial started with a random seed rhythm sampled uniformly from the same 54 ratios of the 2iRT. We presented 3 repetitions of the seed, and the subject needed to synchronize his taps to the repeated rhythm (again, 6 taps for 3 rhythms). At the end of a trial, each inter-response interval (r_x_ ot r_y_) was averaged across the 3 repetitions, and the resulting mean interval was used to form a new seed (Figure 3a). This mechanism was iterated fifteen times. In all cases, an evolution from the original seed distribution was observed, with a convergence towards a multimodal distribution (Figure 3b).

**Figure 3.**
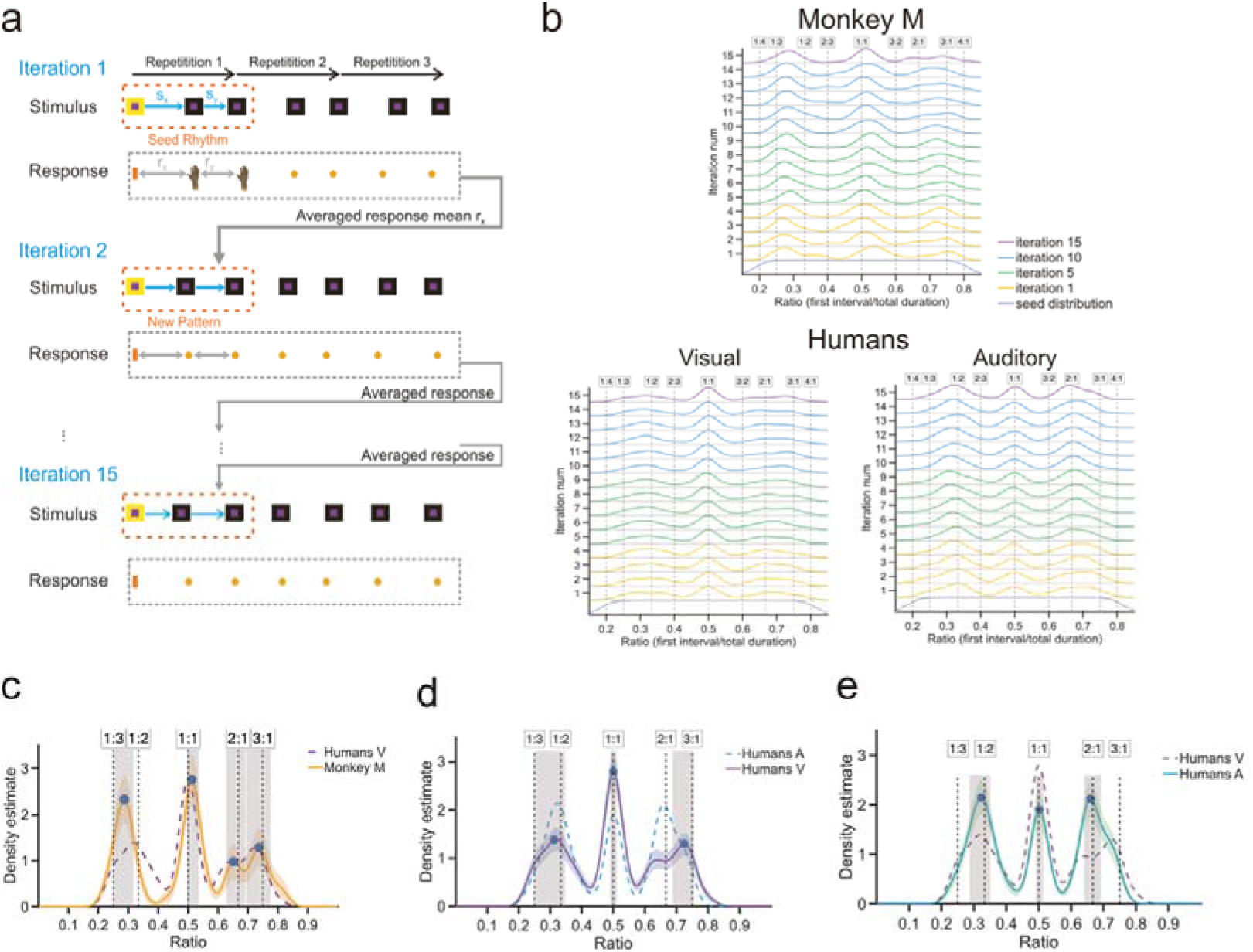
| Iterated Reproduction of Two-interval Beat task. **a,** Structure of the iterated task. The experiment was identical to the two-interval beat synchronization task except that in the testing epoch the monkey performed 15 iterations using the average of the two produced intervals of the previous iteration instead of 15 trials with the same ratio. **b,** Kernel density estimates across iterations for humans (visual n= 25, audio n=14) and monkey M. Aggregated results from the 54 tested ratios by iteration are presented. **c,** Comparison of the distributions of produced ratios for the 15^th^ iteration in both primate species in the visual modality. Monkey distribution corresponds to the yellow line, 95% confidence intervals (CI) were obtained by bootstrap and correspond to the yellow shaded area. The distribution for the human visual condition is shown as the violet dashed line. Statistically significant peaks in the monkey distribution are indicated by the blue dots, and the corresponding 95% CI are shown as the gray shaded areas. Dashed black lines indicate integer ratios. **d**, Similar plot as **c** for the comparison between the two modalities for humans. The violet line is the distribution for human visual. Dashed light blue lines correspond to the human auditory. **e**, Comparison of the human distributions between auditory, light blue line, and visual modality, dashed violet line.

In order to determine the improvement across iterations, we computed the copying accuracy, defined as the Euclidean distance between the target interval and the produced interval. The copying accuracy decreased significantly across three sets of iterations (1-5, 6-10, and 11-15), although the dynamics of reproduction differ between species (see Figure S6a). To further show the accuracy of the reproduction in the iterated task, we computed the difference between the produced ratio and the target ratio for the first and the 15th iteration for monkey M and humans. Biases were overall small for the last iteration for both subjects (Monkey M: mean bias x = 0.0035 CI=[-0.0978, 0.0862]; Humans Visual: mean bias x = 0.0016 CI=[ –0.0990,0.1129]; Humans Auditory: mean bias x = 0.0030 CI=[-0.0661,0.0639]). Additionally, in the final iteration, we identified the preferred categories for the produced ratios (Figure S6b-d).

The continuous distribution underlying the responses was estimated using the kernel density estimation. Figure 3b shows the evolution of these density estimates for each iteration as a function of the target ratio. The distributions rapidly diverged from the seed distribution and converged toward a stable steady state. Specifically, from iteration 3 onward, all iterations exhibited significantly greater distances than expected under the null distribution (all p < 0.002; distances quantified using Jensen–Shannon divergence; see Methods). In contrast, comparisons between iterations 6–10 and 11–15 were non-significant across all three conditions (human audition: JSD = 0.002, p = 0.773; human vision: JSD = 0.0016, p = 0.112; monkey: JSD = 0.0138, p = 0.098), suggesting that the iterative process had largely converged by the 15th iteration.

To identify the preferred categories in the last iteration, we employed a peak-finding algorithm (see methods) over the aggregated produced ratios for the monkey in the visual condition (Figure 3c) and in the visual and auditory conditions of the human (Figure 3d,e). In the three groups, there was a large emerging peak in the isochrony (1:1) with the monkey peaking at 0.51 and a range of 0.50-0.54; while humans in the visual and auditory conditions peaked at 0.50 with a range of 0.49-0.51. Notably, the results from the monkey showed rhythmic categories with integer ratios in the long-short configurations, with peaks at 0.65 [0.63-0.68] and 0.74 [0.70-0.76] corresponding to the 2:1 and 3:1 ratios, respectively (Figure 3c). In contrast, for the short-long configuration, the monkey showed significant peak at a non-integer ratio of 0.29 (∼1:2.5) [0.26-0.32]. The human visual distribution showed two significant peaks around the ratios of 1:2 and 3:1 (0.31 [0.26-0.33] and 0.72 [0.70-0.74]) (Figure 3d), while the human auditory distribution exhibited significant peaks in the expected ratios of 1:2 and 2:1 (0.32 [0.28-0.34] and 0.66 [0.65-0.68]) (Figure 3e).

Finally, to further compare the distributions, we computed the JSD between groups. The difference between the monkey and the human visual distribution was not significant (JSD=0.055, p=0.1) (Figure 3c), whereas the difference between the monkey and the human auditory distribution was highly significant (JSD=0.106, p<0.002). The distance between the two human distributions was slightly significant (JSD=0.042, p=0.022).

Overall, these findings suggest the following. First, macaques and humans exhibit a substantial innate capacity for isochrony. Second, the rhythmic perceptual priors for ratios greater than 0.5 are observed at simple integer ratios in both species. Third, for ratios below 0.5, macaques possess a non-integer rhythm category. Finally, the monkeys’ overall distributional responses resemble those of humans, especially in the visual condition.

## Discussion

Testing the tapping abilities of Rhesus monkeys to synchronize to isochronous and two-interval rhythms we found a series of complex rhythmic skills previously attributed only to humans. First, when properly trained to tap with asynchronies close to zero and to produce intervals close to the metronome’s tempo, macaques show a notorious flexibility to produce intervals with a robust period and phase, across a wide range of isochronous durations. Second, perfect asynchronies to metronomes around 650-850 ms are concordant with the notion of a preferred tempo. Third, monkeys can easily generalize from tapping to isochronous metronomes to produce short-long or long-short interval rhythms. Four, once they learned to perform the 2iRT, the monkeys developed a tempo-invariant representation of ratios, quantifying rhythm in a relative rather than absolute fashion. Fifth, monkeys show categorical rhythmic behavior. Finally, in the iterative task with a full range of ratios, macaques showed similar kernel density distributions as humans in the visual condition, with a large prior for isochrony and integer ratios (2:1, 3:1). Nevertheless, for short-long rhythms monkeys showed a preference for a non-integer ratio of ∼1:2.5, consistent with non-integer ratios observed in some human musical cultures (Polak et al., 2016, 2018; Roeske et al., 2020).

Previous studies have shown that macaques can predictively synchronize to an auditory or visual metronome during synchronization tapping tasks and can also maintain the tempo without the entraining sensory signal in the classical synchronization continuation task(Betancourt et al., 2023; Gámez et al., 2018; Zarco et al., 2009). A key new result of the present study is the wide range of tempos that the monkeys could properly synchronize due to the large emphasis during training to maintain both precise period and phase for each tap with respect to the metronome to receive a reward. Thus, the monkeys developed a predictive and flexible ability to perceive the isochronous tempo and to entrain their tapping. On the other hand, the kinematic analysis of the speed of the tapping hand revealed that monkeys produce a stereotypic movement to push the button across tempos and the serial order elements of the sequence, while the rhythmic timing is controlled on the dwell between movements(Betancourt et al., 2023; Donnet et al., 2014; Gámez et al., 2019). Therefore, these observations suggest that during tapping synchronization, monkeys use a complex rhythmic timing mechanism that includes at least three components. One, a component for controlling the dwell to follow the metronome with different tempos; two, another for triggering the internal beat signal that coincides with the tapping times; and three a component that generates a command that initiates the stereotypic 2-element tapping movement. Preliminary video analysis in the present study showed the same strategy and hand speed profile. Remarkably, specific neural correlates for each of these components have been found in the population activity of medial premotor areas of macaques performing the same tasks. The duration of the dwell is encoded in the amplitude and speed of the neural population trajectories, the internal beat at the tapping times is triggered when the neural trajectories reach a particular zone in state space, and the stereotypic movement is controlled by the speed of the neural trajectories few hundreds of milliseconds before the execution of the up-down and down-up stereotypic movements(Betancourt et al., 2023; Gámez et al., 2019; Lenc et al., 2021). A crucial aspect of these neural population trajectories is that they form elliptic loops of each produced interval in the rhythmic sequence, where time is encoded relatively as the proportion of time that the trajectories have traversed (Betancourt et al., 2023; Gámez et al., 2019). This relative representation of time could be used to encode the observed tempo-invariant ratios during 2iRT. Based on this background we have two specific predictions. One, the neural trajectories will form figure eights of each repetition of the two-interval rhythms, where the proportional geometry of the state space neural population dynamics will encode the ratio. For example, two ellipses with a large and small diameter will encode a long-short rhythm. The second hypothesis is that the priors in the iterative task at isochrony and at the simple ratios of 2:1 and 3:1 will be represented in the precision of the curvature in the elliptical neural trajectories(Remington et al., 2018). If this is true, we could further suggest that the robust priors at simple ratios in humans are due to the geometric properties of the neural trajectories in the medial premotor areas, consistent with recent evidence of neural encoding of rhythm priors in humans (Barbero et al., 2025).

The present findings support the notion that monkeys possess a sophisticated timing mechanism for rhythmic behavior with the ability to produce two interval rhythms over a wide range of ratios and tempos, and with rhythmic priors similar to humans. These results demand a correction in the gradual audiomotor evolution hypothesis (GAE)(Merchant & Honing, 2014). The original GAE suggests that beat-based timing was developed gradually in primates over evolutionary history, with complex rhythmic abilities in humans and basic isochronic skills in monkeys. Here, we showed that macaques, when correctly trained, can develop most of the rhythmic abilities of humans, levering their capabilities from basic isochrony to complex and categorical rhythmic priors using relative timing(Ayala et al., 2017; Criscuolo et al., 2023).This notion has been recently supported by biomusicology studies showing that chimpanzees can learn to execute human-like sequences(Dufour et al., 2015) and produce isochronous drumming on trees (Eleuteri et al., 2025), gibbons produce isochronous songs when dueting, while lemurs produce in the wild coordinated songs with isochronous and 1:2 rhythmic categories(De Gregorio et al., 2021).

In sum, this study provides evidence of a sophisticated rhythmic mechanism present in monkeys and opens the possibility to determine in further experiments the neural underpinnings of complex rhythmic behavior using high-density extracellular recordings of multiple brain areas in the behaving macaque.

## Methods

### Human participants

A total of 25 human participants (13 females and 12 males), mean (SD) age of 30 (±3.76) years, (range: 21-36 years) were tested in this study. They were right-handed, had normal or corrected to normal vision, and were naive about the task and purpose of the experiment. Each subject volunteered and gave informed consent which complied with the Declaration of Helsinki and was approved by the National University of Mexico Institutional Review Board.

### Monkeys

Two Rhesus monkeys (*Macaca mulatta*) referred as monkey M (male, 12.5 kg BW) and monkey D (female, 5.5 kg BW) were used. All the animal care, housing, and training procedures were approved in the protocol 0.81H by bioethics in Research Committee of the Instituto de Neurobiología, Universidad Nacional Autónoma de México The protocol followed the 3Rs and conformed to the principles outlined in the Guide for Care and Use of Laboratory Animals (NIH, publication number 85-23, revised 1985) and the NORMA Oficial Mexicana NOM-062-ZOO-1999, Especificaciones técnicas para la producción, cuidado y uso de los animales de laboratorio’. Monkeys are monitored daily by researchers and the animal care staff to check their conditions of health and welfare.

### Apparatus

Human participants were seated comfortably in a chair facing an 8×8 inch LED matrix (Adafruit NeoPixel, refresh rate of 1000 Hz, 56 cm away from their eyes), placed their right hand on a key with an infrared optical sensor (Balluf BOS 11K, response time ≤ 1 ms), and tapped to a metal button equipped with a proximity sensor (NBB2-12GM50-E2-V1). They performed the tasks in a soundproofed and quiet room. Tucker-Davis Technologies (TDT) hardware (RZ2) and software (Pynapse) were used to deliver stimuli and collect behavioral data at 12kHz. The same setup was used in the monkeys, which were seated in a primate chair while performing the tasks.

### Isochronous synchronization tapping task (ISTT)

The two monkeys were trained in this task. A trial started when the monkey held a lever during the presentation of a yellow background and the first flashing red-square that acted as metronome (Figure 1a). The disappearance of the yellow background was the go-signal to tap a button seven times in synchrony with the red-square metronome, generating seven inter-response intervals (IRI0 to IRI6). The inter-stimulus interval (ISI) of the metronome was randomly selected from 16 intervals (400 – 1150 ms) in blocks of trials. Each block consisted of five training trials, where the monkey adapted to the new tempo, and twenty testing correct trials. The monkeys completed between 10 to 12 blocks in a day. Monkeys received a reward (few drops of water) when two conditions were satisfied: produced intervals in the trial were not different by more than 22% of the instructed interval and all asynchronies between stimuli and taps were less than 22% of the instructed interval. In addition, the reward was doubled or quadrupled when the absolute asynchronies were between 50-100 ms or between 0-100 ms, respectively, to promote predictive timing.

### Two-interval rhythm task (2iRT)

Two monkeys were trained in this task. In each trial, the monkeys produced three repetitions of the two-interval rhythm for a total of six taps. We used 53 ratios (from 0.29 to 0.71) and eight total durations (1,000 to 1,700 ms), for a total of 424 rhythm combinations that were presented pseudorandomly in blocks of trials. Each block was composed of 5 training trials, to get the new rhythm, followed by 15 testing trials. The minimum interval in all combinations was 294 ms and the maximum was 1,200 ms. Importantly, we used color and size cues on the flashing beat stimuli depending on the duration of the target interval: for intervals greater than or equal to 750 ms we used large red squares, and for intervals less than 750 ms we used small green squares. Monkeys performed between 10 to 15 blocks each day. For a block to be considered valid and analyzed further, 40% of the trials needed to be correct (15 trials), otherwise, the block was aborted and that combination of ratio/duration reentered the pseudorandom procedure. Monkeys received a minimum reward when each asynchrony in a trial was less than (±)150 ms. In addition, the monkeys received larger amounts of reward when the mean produced ratio in a trial was close to the target ratio, obeying the following reward rule. Considering the 6 intervals in the 2iRT we have:

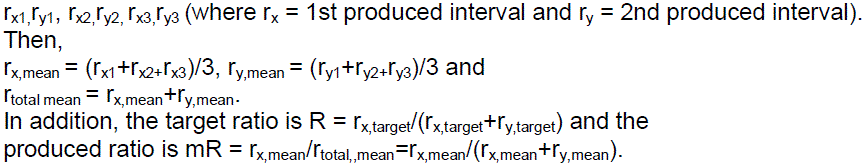

Thus, the reward rule was defined as RR = abs(R – mR) and the amount of liquid reward was given by the values of RR in Table 1.

**Table 1.**
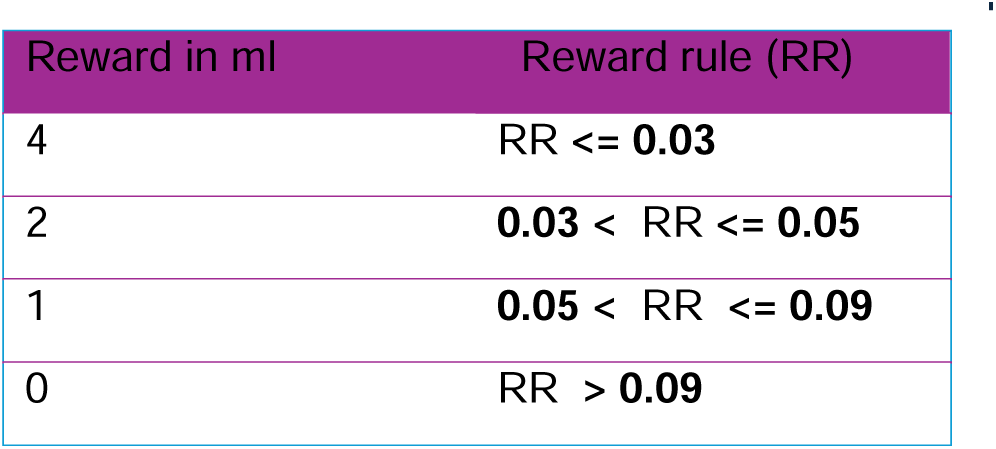
| Reward rule.

| Reward in ml | Reward rule (RR) |
| --- | --- |
| 4 | $RR \leq 0.03$ |
| 2 | $0.03 < RR \leq 0.05$ |
| 1 | $0.05 < RR \leq 0.09$ |
| 0 | $RR > 0.09$ |

### Full-range 2iRT

This experiment was performed in monkey M and we used 54 ratios from a uniform distribution ranging from 0.18 to 0.82. We chose the total duration of 1600 ms since monkeys were quite proficient at producing the two intervals within this period. Again, each ratio was chosen pseudorandomly and each block of trials with a chosen ratio was composed of 5 training trials, to get the new rhythm, followed by 15 testing trials. Notably, for this task the metronome was a uniform flashing square (violet flashing metronome) and hence no short/long cues were provided to the monkey. In addition, the amount of reward for correct trials was always the same. A trial was considered correct when each asynchrony was within a window of (±) 160 ms.

### Iterated task

Humans and Monkey M were tested in this task that had the same ratios, total duration, block structure, and reward contingencies of the full-range 2iRT. The main difference was in the testing epoch of a block. Instead of repeating the same ratio seed for the 15 trials task, the seed of each trial, here iteration, changed depending on the response of the subject. In the first iteration, a seed (a two-interval rhythm) was randomly selected from the 54 range of ratios. From the second to the fifteenth iteration, the target ratio was defined as the average of the IRIs of the previous trial as follows: Each IRI (r_xn_ and r_yn_) of the rhythm was averaged across the 3 repetitions in a trial (mean of r_x_ and mean of r_y_). Then the mean produced ratio was computed, and the new ISI (s_x_) was established for the next iteration. The second ISI (s_y_) was computed as the difference between the total duration of the two-interval rhythm and the s_x._ The new pattern (s_x_ + s_y_) was presented in the next iteration, and this process was repeated 15 times. The monkey completed 810 iterations, humans in the visual modality 4036, and in the auditory modality 1680 iterations.

### Monkey training

Macaques were trained following operant conditioning techniques(Merchant, Crowe, et al., 2011; Naselaris et al., 2006). They received normal food rations, were water-deprived and received most of their water intake during the training and testing sessions. The animals worked 5 days/week, 3 hr/day on average and performed around 200 correct trials per day, with a total liquid intake of 150–300 ml during performance. Weight was strictly controlled; supplementary fluids were provided when monkeys did not reach their minimum water volume on a day of performance.

*ISTT*. Monkeys M and D were initially trained with an isochronous visual metronome for 3 months. First, the monkeys received a reward whenever they held their hand in the lever for a few seconds. Afterward, they learned to push a button after holding the lever. Next, they learned to hold the lever and push the button once a visual stimulus was presented. This first phase lasted 3 weeks. Throughout the following two months, the monkeys were presented with isochronous stimuli with target intervals varying from 550 to 850 ms. Critically, the reward was delivered only on trials with asynchronies close to zero (± 200 ms) with extra rewards for smaller asynchronies. The number of stimuli and required taps in a trial only increased when the monkeys were stable at producing predictive asynchronies. This training strategy was very important for the monkeys to understand the predictive nature of the tapping tasks. Once they reached 70% of correct trials in the full version of the isochronous task (six taps), we collected the data for this paradigm.

*2iRT*. The first days the monkeys confronted the *2iRT* we used a perception epoch at the beginning of each trial to show the rhythm they needed to reproduce. Thus, while macaques held the key, we presented three flashing stimuli defining one repetition of the two-interval rhythm, after which the yellow background disappeared, acting as a go signal to start the reproduction of the beat for three repetitions. From the first day the monkeys were able to reproduce short-long/long-short sequences with surprising flexibility. Nevertheless, they tended to produce the previous instead of the actual target interval (Figure S3). To change this behavioral pattern, we modified different aspects of the original *2iRT.* First, we removed the perceptual epoch and trained the monkeys to start tapping after the first flashing stimulus and the disappearance of the yellow background, as in the original human task (Jacoby and McDermond, 2017). The monkeys, then, needed to recognize the beat pattern, paying attention to the two actual durations. Second, we simplified the trials by using only one repetition of the two-interval rhythm, requiring only three taps, and using intervals with limited temporal differences, ranging only 50 to 250 ms (e.g. 720 – 780 ms, 550 – 800 ms, 840 – 670 ms). Each ratio was presented in blocks of 20 to 50 trials. If the monkeys were struggling with a ratio, we introduced a couple of isochronous blocks to get their performance back, and then continued with the 2iRT. Once the monkeys reached 60% of correct performance, we progressively increased the number of repetitions of the ratio in a trial. During all these stages, the amount of reward was always the same, the goal was to let the monkeys understand the basis of the task. Once the monkeys were stable with three repetitions, we started the training of the 2iRT described above.

### Data analysis

All offline data processing and analysis were performed in MATLAB (2022b, MathWorks). Performance measures (asynchrony, constant error, and Weber fraction) were computed for all the tasks. Asynchronies are defined as the difference between tap onset and stimulus onset. Constant error is the difference between produced and instructed interval. Weber fraction is equivalent to the ratio of the standard deviation of the asynchronies in a trial and the target interval (ISI).

In IST for each performance measure with respect to the sixteen different intervals, we used one-way ANOVA and circular test (Rayleigh test) to evaluate that the different performance measures for each tested interval were significantly different.

In 2iRT we performed t-test, one and two-way ANOVA to evaluate the effect of the total duration and target ratio over the performance measures of interest.

We measured the relationship between target and produced interval or ratio using linear regression function from MATLAB *fitlm*.

For the iterated experiment we followed established procedures used in human experiments (Jacoby et al., 2024; Jacoby & McDermott, 2017).

*Copying accuracy*. We computed copying accuracy as the root mean square difference between the target interval and the produced interval.

We average the responses for each set of iterations (1-5, 6-10, and 11-15) to compute the copying accuracy. We then averaged the result over all the chains by set of iterations.

*Kernel density estimate* (*kde*). To approximate the distribution of preferred integer ratios, we computed the probability density function (PDF) at each iteration using kernel density estimation, leveraging the presence of multiple repetitions that reinforce preferred ratios. A fixed bin size of 0.025 was used for all analysis. To facilitate comparison between distributions, the distribution estimates were normalized relative to the uniform distribution (seed distribution). Namely, if the distribution *p(x)* represents the result of the kernel density, and *u(x)* represents the uniform distribution over the range of possible rhythms (ratios) then the normalized relative to uniform distribution is *p’(x)=p(x)/u(x)*.

*Jensen-Shannon Divergence (JSD) method*. The Jensen-Shannon divergence (JSD), a measure of similarity between two different distributions where 0 is similarity and 1 high difference. To compare two distributions P and Q, the JSD, is defined as follows:

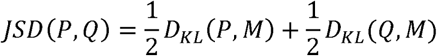

where 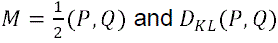 is the Kullback-Leibler divergence:

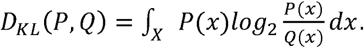

It should be noted that the JSD is symmetric and constrained between 0 and 1, and is equivalent to 0 when *P=Q*.

*Peak finding.* To identify statistically reliable peaks in the regions corresponding to simple integer ratios (e.g. 1:3, 1:2, 1:1…) from iterated data, we created 1,000 bootstrapped datasets by sampling the original collected data with replacement over the blocks of testing stage of the task; for each bootstrap dataset, we computed the kernel density estimate associated with the data (we used a bandwidth of 0.025 in all analysis), then we computed all peak locations for a given bootstrapped dataset using MATLAB’s *findpeaks* function with default parameters. To determine the average number of peaks detected, we count the number of bootstraps contained in a region of interest defined by the different integer ratios +/− 0.033, after that we selected the regions that were present in more of the 80% of bootstrapped datasets.

*Significance of the difference between seed distribution and iterations*. To obtain statistical significance of the difference between the two distributions, we used bootstrapping. For each iteration, we applied kernel density estimation to these samples and computed the JSD and compared it for the same distance computed between the real seeds and a sample with the same number of trials but sampled from the seed distribution. We repeated this process 1,000 times to obtain significance level.

*Significance of the difference between sets of iterations*. To compute the significance of the JSD between two sets of iterations (for example data from a Monkey and a group of humans), we permuted the chains between the two sets to compute the corresponding JSD. Namely, we reshuffled all trials into two new randomized groups of the same size. We then applied kernel density estimation to these samples and computed the corresponding JSD. We repeated this procedure 1000 times to compare the actual JSD with the resulting null distribution and with that obtain the probability of the actual JSD.

## Acknowledgments

We thank Zayani Medrano, Maria Antonieta Carbajo, Juan Ortíz, and Raul Paulín for their valuable technical assistance. Supported by CONACYT: A1-S-8430, PAPIIT: IG200424, and UNAM-DGAPA-PASPA.

Ameyaltzin Castillo-Almazán is a doctoral student from the Programa de Doctorado en Ciencias Biomédicas, Universidad Nacional Autónoma de México (UNAM) and has received a fellowship (No. 963683) from Secretaría de Ciencias, Humanidades, Tecnología e Innovación (SECIHTI, formerly CONAHCYT).

## Author contributions

H.M., N.J. and A.C-A. conceived the experiments. A.C-A. and L.P. collected psychophysics data. A.C-A. analyzed the data. H.M. and N.J. supervised the project.

