## Supplementary Figures for "Categorical rhythmic priors in macaques"

### Supplementary information

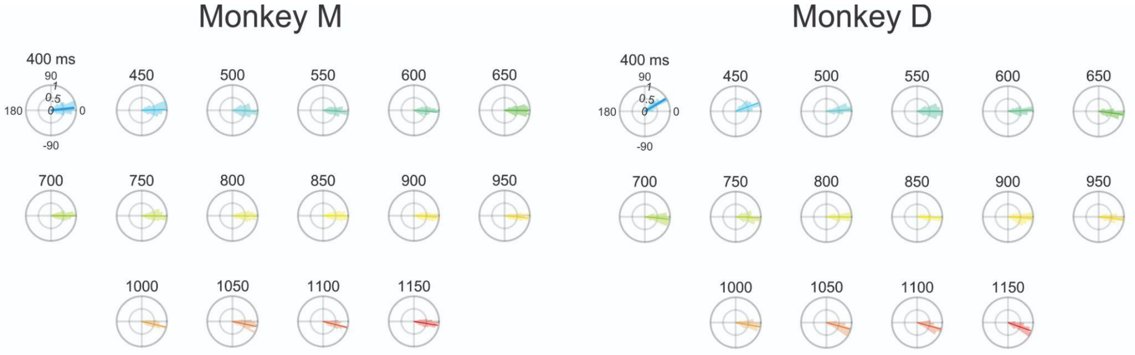


**Figure S1 |** Circular plots of mean asynchronies for the whole range of tested intervals for both monkeys. Asynchronies for both monkeys are not uniformly distributed (Rayleigh test p<0.000001, monkey M z = 5.82e3; monkey D z = 2.11e3) and the corresponding circular t-test (circ_mtest in matlab) showed that both monkeys have a preferred ratio around 750 - 850ms where the asynchronies were not different from zero.

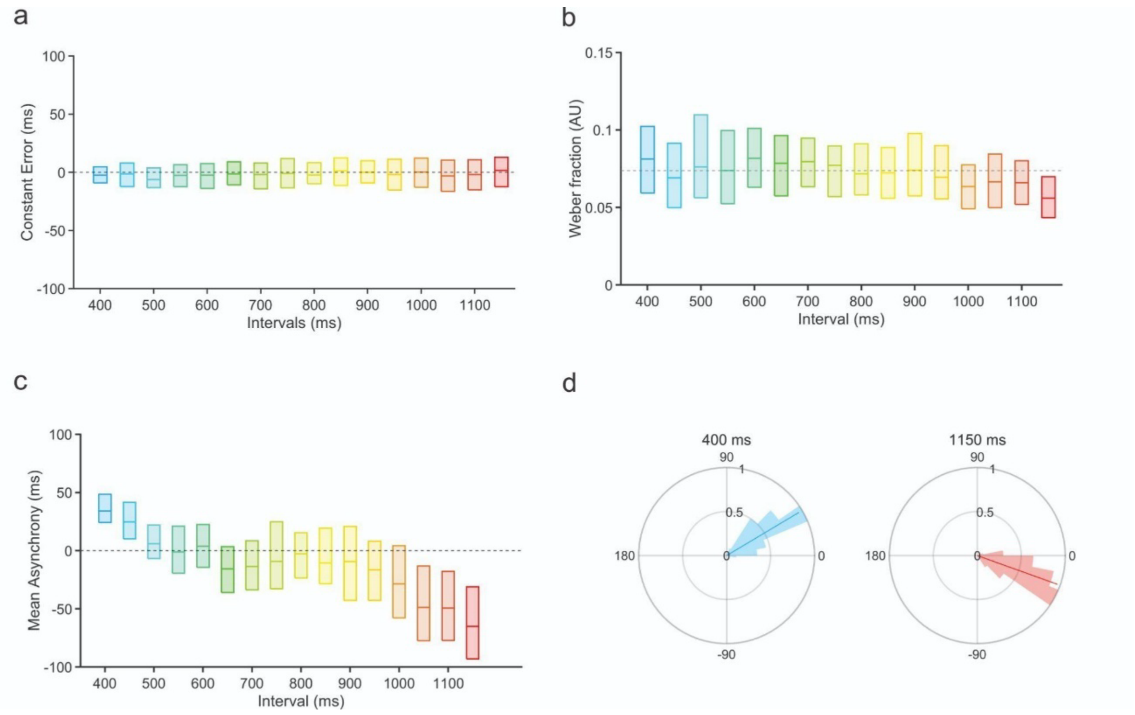


**Figure S2 |** Isochronous tapping synchronization task and behavior for monkey D. **a**, Boxplots of the Constant Error (CE) as a function of interval showing the median (central line), the 25th (bottom edge) and 75th percentiles (top edge). There was no statistically significant difference between the mean CE for the different intervals, suggesting that all the different intervals were precisely reproduced. (One-way ANOVA: $F(15,2399) = 1.23$, $p = 0.24$ $\eta^2 = 0.008$) **b**, Boxplot of Weber fraction for each tested interval. The monkey shows a lower Weber fraction which witnesses a high uniform performance for all intervals, even though there is a difference between intervals (One-way ANOVA: $F(15,2399) = 10.05$, $p = 1.18e^{-23}$ $\eta^2 = 0.059$). **c**, Asynchrony Boxplots. The monkey's taps were predictive, with asynchronies close to zero across the wide range of tested intervals. Nevertheless, there was a significant difference between the mean asynchrony as a function of interval (ANOVA, $F(15,2399) = 76.64$, $p < 7.11e^{-191}$ $\eta^2 = 0.325$), mainly due to the negative asynchronies for long intervals. **d**, Polar plots showing examples of the distributions of the asynchronies for the extreme tested intervals: shorter 400 ms and longer 1150 ms. We found that the mean resultant was close to one (showed as a vector; 0.96 and 0.97, respectively) indicating large tapping consistency, with a significant unimodal distribution (both intervals with a significant Rayleigh's test $p < 0.00001$), and that displayed an angle that was close to zero, reflecting large rhythmic tapping prediction.

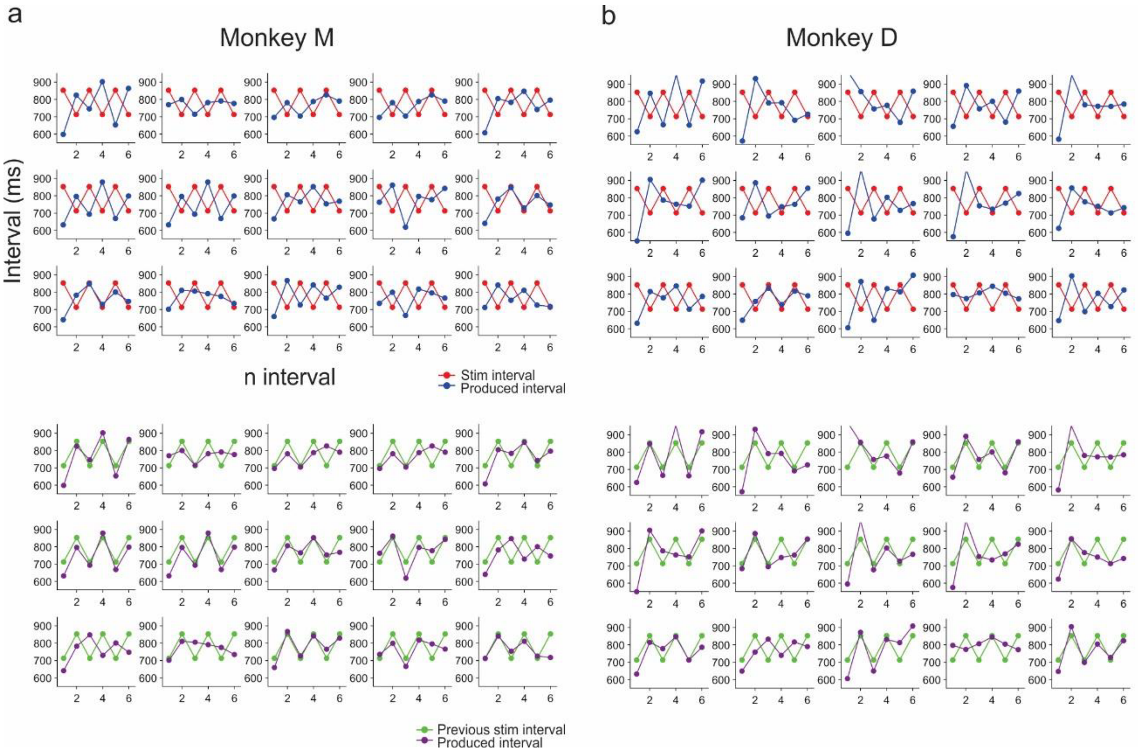


**Figure S3 |** Examples of produced intervals by trial for 2iRT. **a**, Example for a block of the rhythm 680 – 820 ms the first day we tested the 2iRT on monkey M. Above, we have 15 trials with the currently displayed interval (red) and the currently produced interval (blue). Below, the same 15 trials are presented with the produced interval (purple) aligned with the previously presented interval (green). **b**, The same rhythm is shown for monkey D, and also for the first day the monkey did the task.

a

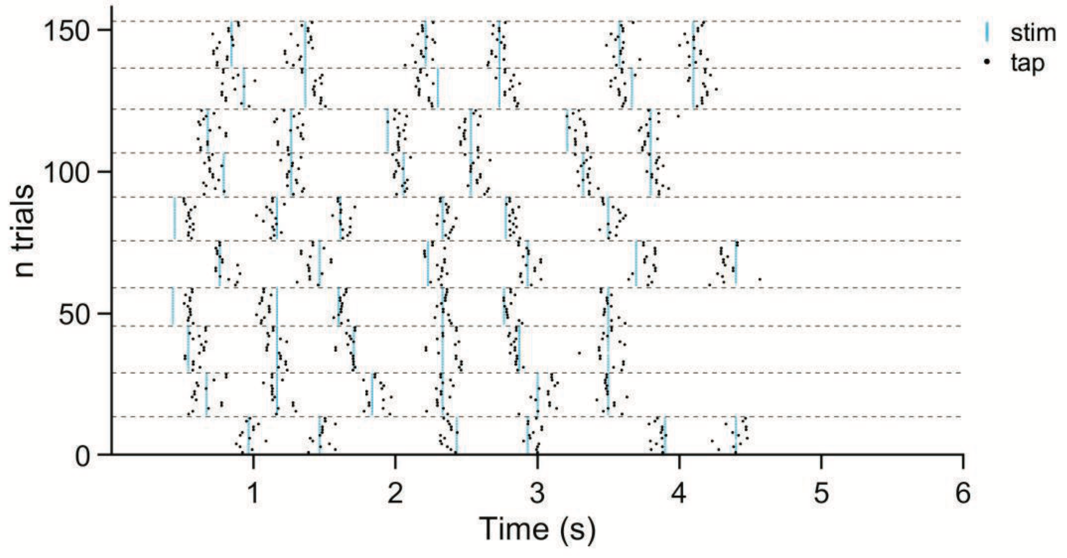


b

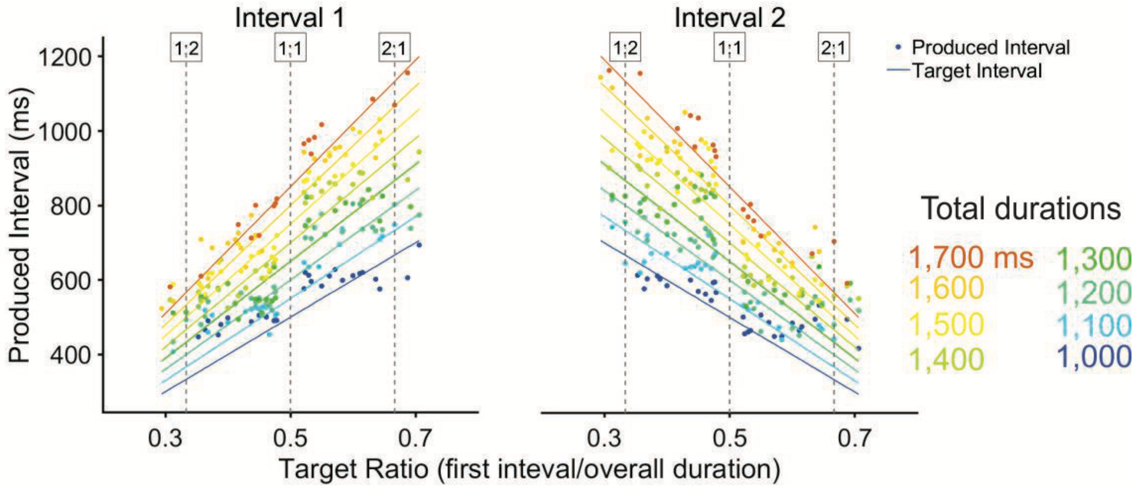


**Figure S4**. | Two-interval rhythm task for monkey D. **a**, Raster plot for a day of performance in the two-interval beat task. Each block has 20 trials of the same combination of interval ratio and total duration. Each block is delimited by the horizontal dashed gray line. Black dots correspond to taps and blue lines to each presented stimulus. Note the close relation between tapping times to each stimulus of the rhythm. **b**, Mean produced intervals. Dashed gray lines indicate integer ratios on the top. The monkey was able to flexibly change the duration of his two produced intervals depending on the total beat duration and the configuration of the rhythm (fast-slow or slow-fast).

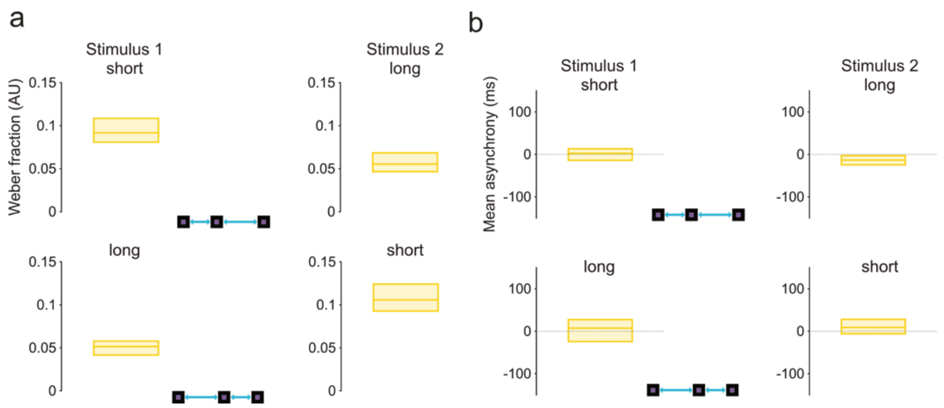


**Figure S5. |** Results for 2iRT full range. **a**, Weber fraction for each stimulus of the two-interval rhythm. **b**, Mean asynchrony. The monkey has very little mean asynchronies for all the ratios, these are close to zero, and generally negative showing his predictive ability.

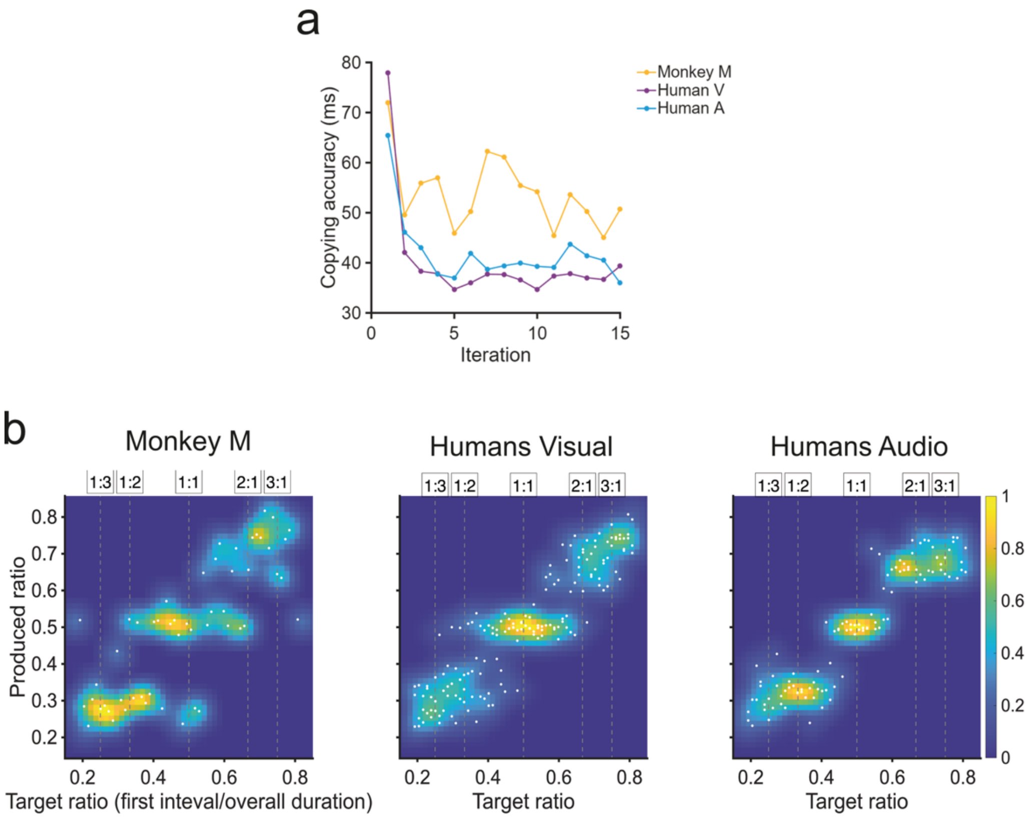


**Figure S6. | Copying accuracy and 2D density plot (related to Iterated task). a,** Copying accuracy by iteration. We compare sets of iterations (1-5, 6-10, and 11-15) with each other. For humans, copying accuracy was significantly different between the first five iterations and the two other iteration sets ($p<0.001$, Audio: $t(107)=2.96$ and $t(107)=2.75$; Visual: $t(161)=6.31$ and $t(161)=4.76$). In addition, no significant effect were observed between the two last sets of iterations (6-10 vs 11-15, Audio: $p=0.85, t(107) = -0.19$; Visual: $p=0.45$, $t(161)=-0.76$). For the monkey, there was a significant difference between the two sets of iterations (6-10 vs 11-15, $p<0.040$, $t(52) = 2.10$). These results suggest that humans had a convergence in the first 5 iterations independently of the synchronization modality. In contrast, the monkey reached this convergence at a later stage. **b**, 2D distribution plot of the produced ratios as a function of the target ratio for the last iteration in the three groups. The preferred rhythmic categories are shown by the heatmap, yellow areas represent high density. The white dots correspond to the produced ratios for the group.
